# *Drosophila* Asap regulates cellular protrusions via dArf6-dependent actin regulatory pathway

**DOI:** 10.1101/2024.01.17.575521

**Authors:** Shikha Kushwaha, Bhagaban Mallik, Zeeshan Mushtaq, Anjali Bisht, Vimlesh Kumar

## Abstract

Membrane protrusions are fundamental to cellular functions like migration, adhesion, and communication and depend upon the dynamic reorganization of the cytoskeleton. The GAP-dependent GTP hydrolysis of Arf proteins regulates actin-dependent membrane remodeling. Here, we show that the dAsap regulates membrane protrusions in S2R+ cells by a mechanism that critically relies on its ArfGAP domain and re-localization of actin regulators, SCAR, and Ena. While our data reinforce the preference of dAsap for Arf1 GTP hydrolysis *in vitro*, we demonstrate that induction of membrane protrusions in S2R+ cells depends on Arf6 inactivation. This study furthers our understanding of how dAsap-dependent GTP hydrolysis maintains a balance between active and inactive states of dArf6 to regulate cell shape.

## Introduction

Eukaryotic cells possess a highly intricate membrane system characterized by the formation of diverse cellular structures such as protrusions, invaginations, and vesicles (McMahon and Boucrot, 2015). These structures play integral roles in fundamental biological processes such as cell division, migration, secretion, fission, fusion, and intracellular trafficking (Kessels and Qualmann, 2015; Liu et al., 2015; Safari and Suetsugu, 2012). The orchestration of these biological phenomena hinges on the dynamic interplay between membrane dynamics and the underlying cytoskeleton, primarily composed of actin and microtubule networks (Mallik and Kumar, 2018; Safari and Suetsugu, 2012; Stanishneva-Konovalova et al., 2016). Several proteins, including Bin/Amphiphysin/Rvs161/167 (BAR) domain proteins, are recognized as critical modulators for inducing membrane dynamics (Mallik et al., 2022; Oh and Robinson, 2012; Richnau et al., 2004). The BAR domain, known to form functional dimers, binds to negatively charged membranes, generating curvature crucial for various cellular events (Peter et al., 2004). Moreover, BAR family proteins exhibit diverse effects on membrane morphology and are further classified into N-BAR, F-BAR (Fes-Cip4 homology BAR), and I-BAR (inverse BAR) classes based on structural properties. While most N-BAR and F-BAR proteins generate tubules, I-BAR proteins induce protrusions upon overexpression (Oh and Robinson, 2012; Safari and Suetsugu, 2012). Nevertheless, exceptions to this generalization, such as the F-BAR protein srGAP2 forming protrusions, underscore the complexity of these regulatory mechanisms (Guerrier et al., 2009).

Besides, BAR domain-containing proteins also interact with small GTPases like Rho and Arf family proteins, further intertwining membrane dynamics with cellular signaling pathways (Coyle et al., 2004; Johnson et al., 2011; Rodal et al., 2008; Rodrigues et al., 2016). One such class, ArfGTPases, a subset of the Ras superfamily, actively participate in membrane trafficking and actin cytoskeleton modulation (D’Souza-Schorey and Chavrier, 2006; Donaldson and Jackson, 2011; Jackson and Bouvet, 2014). In *Drosophila* melanogaster, the single Arf orthologue of each class, namely dArf1, dArf4, and dArf6, governs these processes (Donaldson and Jackson, 2011). However, the role of Arf6 is particularly implicated in plasma membrane endocytosis, exocytosis, endosomal recycling, cytokinesis, and actin cytoskeleton reorganization by ArfGEFs and ArfGAPs (Jackson and Bouvet, 2014). Interestingly, the subfamily of ArfGAPs known as ASAPs is characterized by an N-terminus BAR domain, PH domain, ArfGAP domain, Ankyrin repeats, and SH3 domain (Inoue and Randazzo, 2007).

In mammals, ASAP1, ASAP2, and ASAP3 are reportedly involved in invasion and cancer metastasis, with ASAP1 regulating focal adhesions, invadopodia, and podosome formation in cells (Muller et al., 2010; Oda et al., 2003). In *Drosophila* melanogaster, the ASAP family is represented by a single orthologue, dAsap (CG30372; FBgn0050372), sharing 38.9% identity with human ASAP1 and a similar domain organization. Further analysis revealed that dAsap is implicated in regulating the actin cytoskeleton during pupal ommatidia formation, Golgi function at the cleavage furrow biosynthesis, and embryonic ectoderm plasma membrane localization during the development of the organisms (Johnson *et al*., 2011; Rodrigues *et al*., 2016; Shao et al., 2010). Despite these findings, the underlying mechanisms by which dAsap contributes to membrane dynamics and actin cytoskeleton regulation remain poorly understood.

This study aims to bridge this gap by characterizing the role of dAsap in actin cytoskeleton reorganization during protrusion formation in *Drosophila* S2R+ cells. Our findings demonstrate that dAsap overexpression induces peripheral protrusions, with dAsap co-localizing with the actin cytoskeleton. We reveal that the ArfGAP activity of dAsap is indispensable for protrusion formation. Moreover, our results indicate that dAsap acts as a GTPase-activating protein (GAP) for Arf1 and Arf6 *in vitro*, highlighting its role in regulating actin cytoskeleton reorganization during peripheral protrusion formation in an Arf6-dependent manner. This comprehensive exploration of the dAsap function sheds light on the intricate interplay between membrane dynamics, actin cytoskeleton regulation, and small GTPases, contributing to a deeper understanding of cellular protrusions in eukaryotic cells.

## Results

### dAsap overexpression induces cellular protrusion in S2R^+^ cells

The S2R+ cells vary in cellular architecture and typically exhibit diverse morphology characterized by broad and thin lamellipodia (Biyasheva et al., 2004; Mallik et al., 2023a). ASAP1 has been shown to induce diverse actin-rich structures, including actin stress fibers, dorsal membrane ruffles, focal adhesions, and invadopodia (Gasilina et al., 2019; Muller *et al*., 2010; Oda *et al*., 2003; Randazzo et al., 2000). To elucidate the role of dAsap in actin cytoskeleton remodeling-dependent membrane protrusions, we first reconfirmed the prior findings by overexpressing dAsap (Actin5C-Gal4; UAS-dAsap) in S2R+ cells. In line with previous findings, the overexpression of dAsap led to membrane protrusions in S2R+ cells, which were largely absent in controls. (Figure 1A-F). Next, to highlight the role of Asap and cytoskeleton in lamellipodia formation, we co-stained the transfected S2R+ cells for dAsap (anti-HA antibodies) and Phalloidin (Figure 1A-F) or dAsap (anti-HA antibodies) and Tubulin (Figure 1G-L). Interestingly, the overexpression of Asap primarily resulted in the re-localization of actin and tubulin to filopodia-like structures. The overexpressing cells displayed a major portion of actin but not tubulin relocalized to filopodia, which is otherwise predominantly localized to cytosol and membrane in controls. These filopodia-like structures were also rich in Asap, which co-localized with actin and, in some cases, tubulin. These observations suggest that Asap induces filopodia formation via relocalizing actin regulatory proteins.

**Figure 1:**
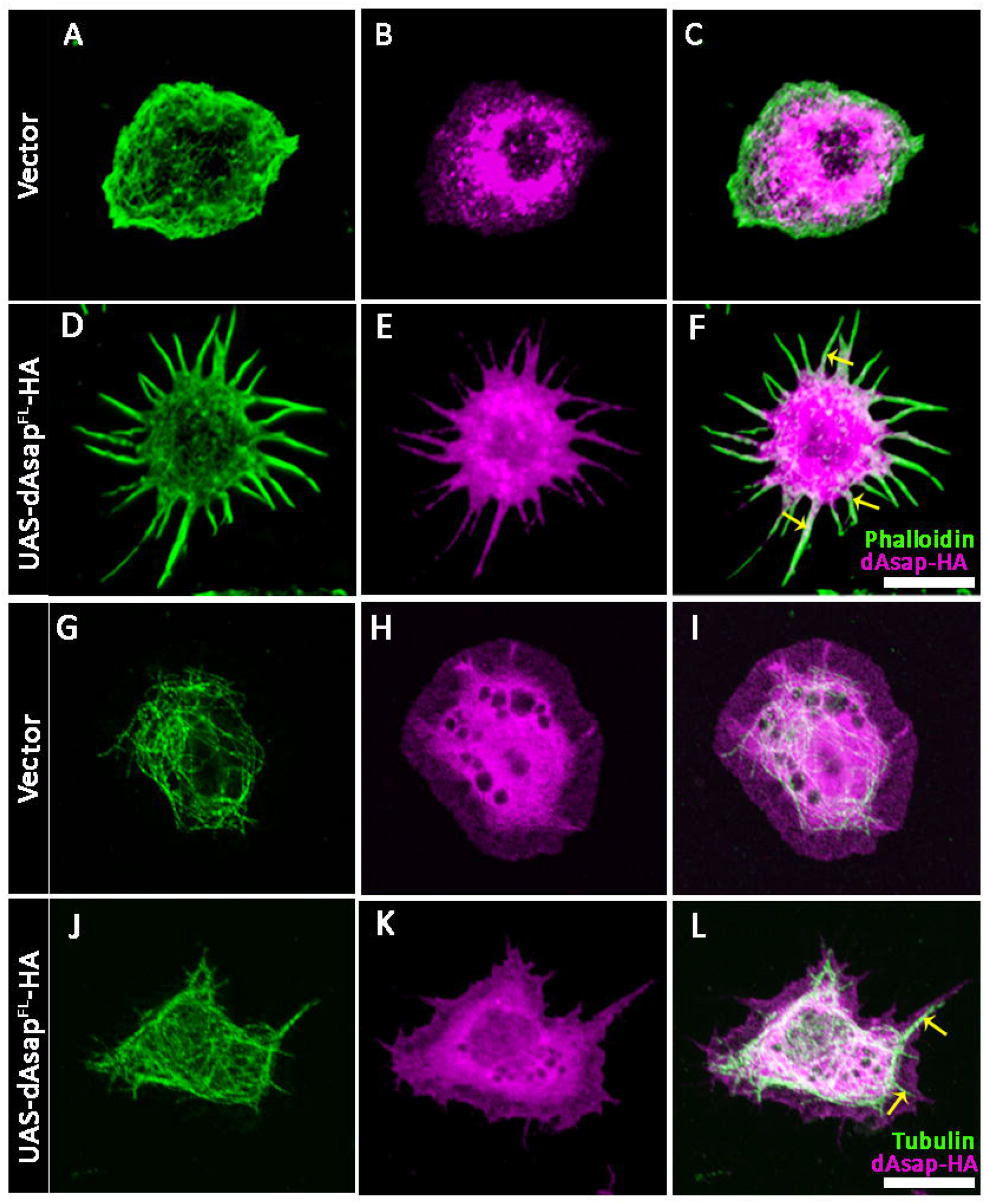
Elevated expression of dAsap leads to the induction of filopodia formation in S2R+ cells. **(A-F)** Representative confocal images show S2R^+^ cells were transfected with control pAWH and HA-dAsap. **(D-F)** Overexpression of dAsap resulted in filopodia formation in S2R^+^ cells compared to empty vector. S2R^+^ cells adhered on Concanavalin-A coated coverslips and immunostained with anti-HA (magenta) and phalloidin conjugated with Alexa-488 (green). HA-dAsap was found to be localized with the actin-rich peripheral protrusions marked with phalloidin. **(G-L)** Overexpression of dAsap did not result in any significant change in the morphology of the microtubule cytoskeleton compared to the empty vector. However, HA-dAsap was found to be co-localized with tubulin, mainly in long filopodia. Scale bar 10 µm.

### The dAsap-Arf-GAP domain is indispensable for generating filopodia in S2R+ cells

*Drosophila* Asap, a homolog of mammalian ASAP1, shares a highly conserved domain organization with ASAP1 (Figure 2A). To test the involvement of each domain in actin cytoskeleton-assisted filopodia formation, we individually overexpressed different dAsap constructs containing different domain(s) in S2R+ cells. Consistently, cells overexpressing dAsap led to the formation of elongated protrusions (dAsap^FL^: # of filopodia, 7.66 ± 0.87, n=110, length in µm: 8.73 ± 0.28, n=67) compared to controls (vector: # of filopodia, 0.67 ± 0.12, n=105, length in µm: 0.42 ± 0.18, n=62) (Figure 2-B-C, 2I-J). Surprisingly, in contrast to our previous findings (Mallik et al., 2023b) and work from others (Ahmed et al., 2010; Krugmann et al., 2001), overexpression of the dAsap^BAR^ domain alone in S2R+ cells did not induce cell protrusions or tubules (dAsap^BAR^: # of filopodia, 0.94 ± 0.14, n=100, length in µm: 0.53 ± 0.21, n=70). These data were comparable to the empty vector controls (vector: # of filopodia, 0.67 ± 0.12, n=105, length in µm: 0.42 ± 0.18, n=62)) (Figure 2D, 2I-J). Interestingly, overexpression of the dAsap constructs lacking BAR domain (dAsap^ΔBAR^: # of filopodia, 7.22 ± 0.91, n=100, length in µm: 7.94 ± 0.20, n=128), induced filopodia similar to full-length dAsap^FL^ (Figure 2E, 2I-J). Further molecular characterization of dAsap showed that dAsap^BPZ^ construct, carrying the Arf-GAP domain, was sufficient to generate a significant number of filopodia in S2R+ cells (dAsap^BPZ^: # of filopodia, 6.37 ± 0.88, n=97, length in µm: 7.80 ± 0.22, n=86) (Figure 2F, 2I-J). Consistently, constructs lacking the Arf-GAP domain, dAsap^ΔPZ^ and dAsap^ΔZ^ (dAsap^ΔPZ^: # of filopodia, 2.16 ± 0.27, n=95, length in µm: 0.72 ± 0.24, n=61; dAsap^ΔZ^: # of filopodia, 1.39 ± 0.24, n=95, length of filopodia: 0.48 ± 0.21), showed a significant but not complete reduction of filopodia formation (Figure 2G-J).

**Figure 2:**
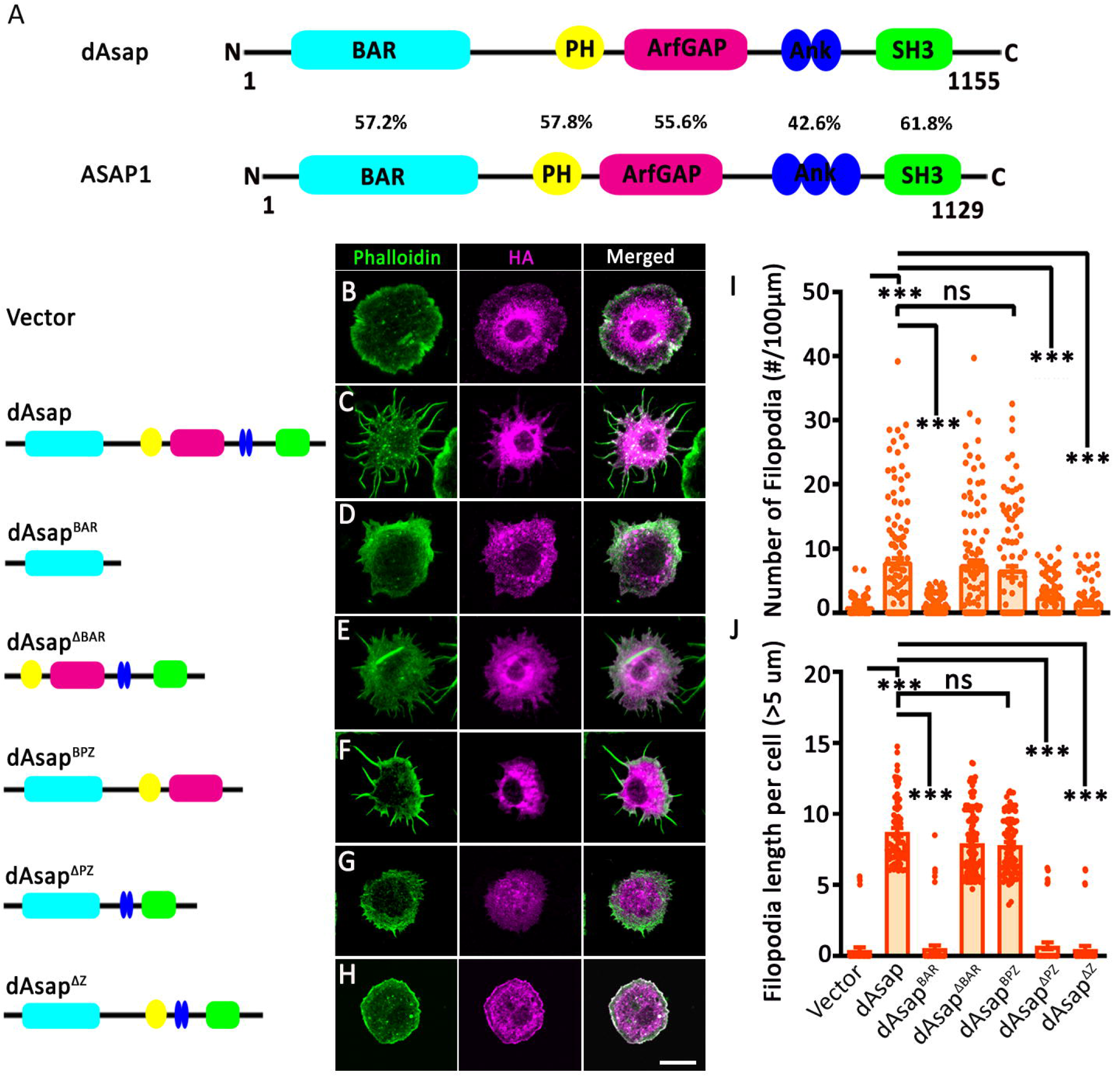
ArfGAP domain of dAsap is crucial for generating filopodia in S2R^+^ cells. **(A)** *Drosophila* Asap is a multidomain protein consisting of N-terminal BAR, PH, ArfGAP, two Ank repeats and an SH3 domain at its C-terminus. Alignment of dAsap shows highly conserved domains and 38.9% overall identity with ASAP1. *Drosophila* Asap consists of highly conserved domains, having its BAR domain-57.2%, PH domain-57.8%, ArfGAP domain-55.6%, Ankyrin repeats-42.6% and SH3 domain-61.8% identical to respective domains of ASAP1 protein. **(B-H)** Representative images of transfected S2R^+^ cells with empty vector and with C terminal HA tag constructs of dAsap. **(D-E)** The dAsap^BAR^ alone does not form filopodia in S2R^+^ cells, and deletion of the BAR domain (dAsap^ΔBAR^) did not inhibit the generation of filopodia. **(F)** However, dAsap^BPZ^ (Z denotes Arf-GAP domain) shows a significant increase in protrusion formation compared to dAsap^BAR^. **(G-H)** The overexpression of dAsap^PZ^ and dAsap^ΔZ^ resulted in the reduction of protrusion number, compared to dAsap **(I-J)** The graph quantifies the number and length of filopodia per hundred-micrometer cell circumference. Actin-rich structures were immunostained with phalloidin, and the expression of different dAsap constructs was marked with anti-HA antibody. Scale bar 10 µm. All values indicate mean and standard error mean. ***p<0.0001, and ns, not significant. One-way ANOVA followed by post-hoc Tukey’s test was used for statistical analysis. Δ denotes the deletion of the particular domain from the full-length protein. Each data point or n represents an individual cell used for quantification.

To discern whether the observed protrusions were lamellipodial or filopodial structures, we co-stained cells overexpressing full-length dAsap with HA (for Asap) and SCAR (a lamellipodia marker) or Ena (a filopodia marker) (Barzik et al., 2014; Hahne et al., 2001; Law et al., 2013; Winkelman et al., 2014). In control cells, both SCAR and Ena appeared as punctate structures localizing majorly to the leading edge of the membrane (Figure 3A-A′, 3C-C′). In contrast, overexpression of dAsap resulted in the redistribution of SCAR and Ena. SCAR showed a more diffused pattern throughout the cell and relocalized to membrane protrusions; however, Ena retained specific wildtype localizations while relocalizing to membrane protrusions (Figure 3B-B′, 3D-D′). In addition, Western blot analysis revealed no significant change in total protein levels of SCAR and Ena in dAsap overexpressing cells compared to controls (Figure 3E). Together, these data underscore the dispensability of the BAR domain for membrane tubule generation and filopodia or protrusion formation in cultured cells. Further, our findings emphasize the critical role of the Arf-GAP domain in actin cytoskeleton-mediated protrusion formation that is assisted to a certain degree by PH domain. Arf-GAP domain induces these lamellipodia through actin cytoskeleton reorganization by relocalizing the actin regulatory proteins, SCAR, and Ena.

**Figure 3:**
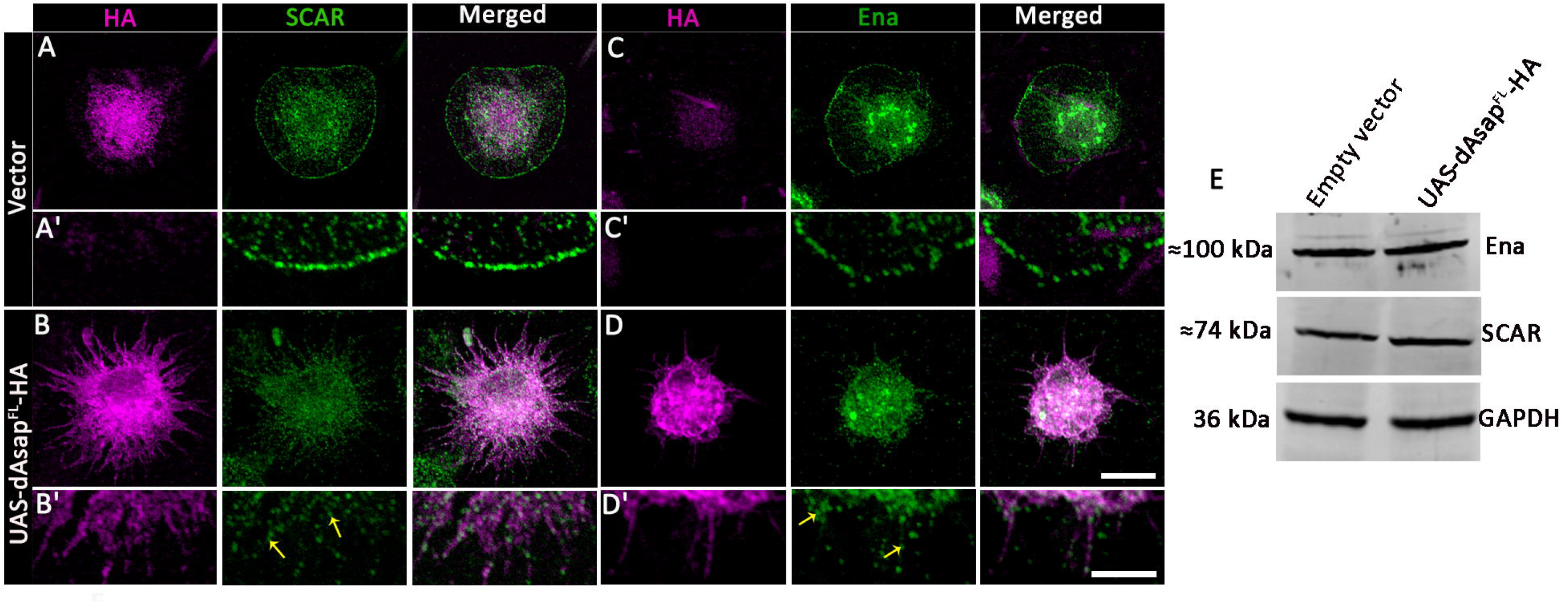
Actin regulatory proteins are relocalized in S2R+ cells following the overexpression of dAsap. **(A-A’, C-C’)** Representative confocal images show SCAR localization in the empty vector and S2R^+^ cells expressing dAsap. Panel **A-A’** represents the localization of SCAR (green) at the cell periphery in S2R^+^ cells; however, in panel **B**, SCAR is redistributed to the filopodial structures in the S2R^+^ cells expressing dAsap (magenta) probed with anti-HA antibody. Panels A’ and B’ show the magnified view of SCAR localized at the cell periphery and then redistributed to the filopodia upon overexpression of dAsap. **(C, C’, D and D’)** Similarly, in panel C, Ena (green) is localized at the cell circumference in cells transfected with an empty vector. In contrast, panel **D** shows the distribution of Ena to the filopodia formed upon dAsap overexpression in S2R^+^ cells. Panels C**’** and D**’** represent the magnified view of localization of Ena in cells expressing empty vector and dAsap, respectively. The scale bar is 10 µm and 3 µm, respectively. **(E)**. Western blot analysis showed that levels of SCAR and Ena were not altered upon overexpression of dAsap in S2R^+^ cells compared to cells transfected with an empty vector. GAPDH was used as a loading control.

### dAsap does not rely on Arf1 to induce filopodia

How might dAsap regulate the actin cytoskeleton to induce filopodia-like protrusions? Mammalian ASAP1 has been shown to regulate actin cytoskeleton via its Arf-GTPase activity (Kam et al., 2000). To test whether the dAsap-Arf-GAP domain is functionally conserved to ASAP1, we analyzed dAsap-Arf-Gap GAP hydrolyzing activity for Arf1 and Arf6. Interestingly, our analysis revealed dArf1-GTP, compared to dArf6-GTP, as the preferred substrate for GTP hydrolysis by dAsap-Arf-GAP (Figure 4A-B). Next, we explored whether dAsap induces membrane protrusion by regulating Arf activity. Owing to the preference of dAsap for Arf1 as a substrate, we first tested the requirement of Arf1 for protrusion formation by overexpressing full-length, dominant negative, or constitutively active forms of dArf1 in cells overexpressing dAsap. Expectedly, compared to control cells expressing dAsap^FL^ led to the formation of protrusions (vector: # of filopodia, 0.35 ± 0.10, n=51;dAsap^FL^: # of filopodia, 15.17 ± 1.35, n=46). In contrast, cells individually overexpressing Arf1^FL^, dArf1^CA^ or dArf1^DN^ did not show any significant change in protrusion formation (Arf1^FL^: # of filopodia, 0.62 ± 0.17, n=41; Arf1^CA^: # of filopodia, 0.60 ± 0.22, n=36; Arf1^DN^: # of filopodia, 0.42 ± 0.11, n=51) (Figure 4 C-H). Interestingly, overexpression of either full-length dArf1^FL^ (dAsap^FL^ + dArf1^FL^: # of filopodia, 14.06 ± 1.56, n=41), a constitutively active form of dArf1 (dAsap^FL^ + dArf1^CA^: # of filopodia, 12.83 ± 2.21, n=37), or dominant negative form of dArf1 (dAsap^FL^ + dArf1^CA^: # of filopodia, 11.98 ± 1.38, n=49) showed no further change in the number of filopodia in cells overexpressing dAsap (Figure 4H). Hence, this data demonstrated that dAsap acts as a GAP for Arf1 and Arf6 but acts independently of Arf1 to induce filopodia.

**Figure 4:**
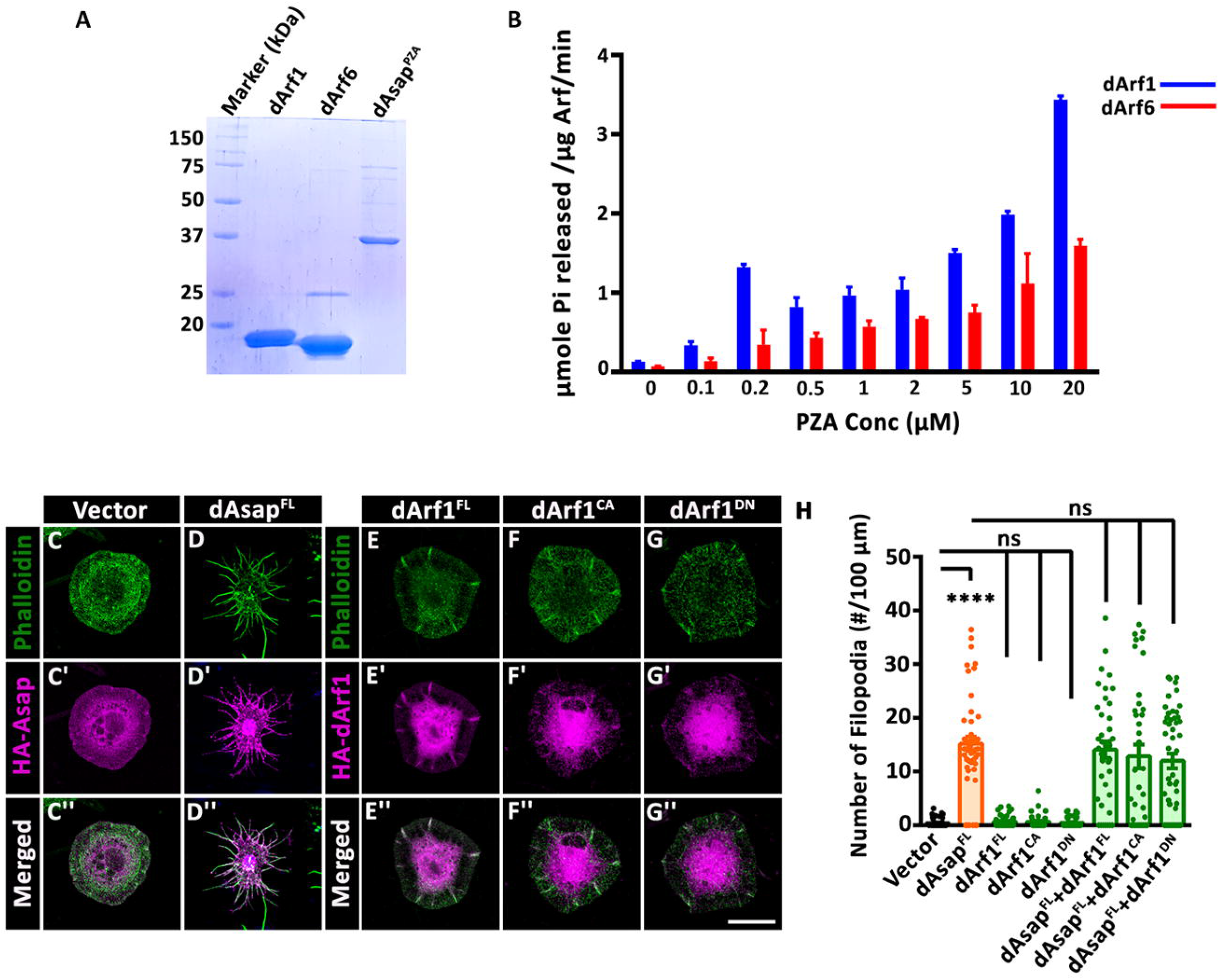
Elevated expression of dArf1 does not mitigate the cellular protrusion induced by dAsap in S2R+ cells. **(A)** Representative gel image stained with Coomassie brilliant blue showing purified proteins, Arf1, Arf6, and dAsap^PZA^ used for the biochemical assay. **(B)** The PZA domain of dAsap has more affinity for Arf1 than Arf6 to hydrolyze the GTP. Arf1 loaded with GTP shows more inorganic phosphate release with increasing concentration of PZA. **(C, C’’- G, G’’)** Representative images of transfected S2R^+^ cells with empty vector, C-terminal HA tag constructs of dAsap and dArf1. **(C, C’’- D, D’’)** The overexpression of dAsap resulted in numerous filopodia in S2R^+^ cells compared to the empty vector. **(E, E’’- G, G’’)** The overexpression of dArf1^FL^, dArf1^DN,^ or dArf1^CA^ resulted in less protrusion formation than dAsap in S2R^+^ cells. Co-expressing dAsap with any of the dArf1 constructs did not suppress the overgrowth of cellular protrusions in S2R+ cells. S2R^+^ cells were immunostained with phalloidin (green) and anti-HA (magenta) antibodies. Scale bar 10 µm. **(H)** The graph quantifies the number of filopodia per hundred-micrometer square of cells. All values here indicate mean and standard error mean. ***p<0.0001 and ns, not significant. One-way ANOVA followed by post-hoc Tukey’s test was used for statistical analysis. Each data point or n represents an individual cell used for quantification.

### dAsap modulates dArf6 activity to induce filopodia in S2R+ cells

Next, we tested whether dAsap regulates Arf6 activity to induce filopodia. Similar to our Arf1 approach, we co-transfected S2R+ cells with HA-dAsap and Myc-dArf6. As expected, dAsap overexpression resulted in a significant enhancement in protrusion number and length. Interestingly, in contrast to dArf1, overexpression of dArf6 led to protrusion formation similar to dAsap (vector: # of filopodia, 1.39 ± 0.20, n=115, length in µm: 0.55 ± 0.21, n=55; HA-dAsap^FL^: # of filopodia, 13.19 ± 0.91, n=117, length in µm: 8.00 ± 0.24, n=143) or Myc-dArf6 (Myc-dArf6^FL^: # of filopodia, 11.81 ± 0.94, n=118, length in µm: 7.20 ± 0.20, n=77). The overexpression of constitutively active Arf6 did not induce filopodia; however, overexpression of dominant negative Arf6 (dArf6^DN^: # of filopodia, 7.65 ± 0.63, n=130, length in µm: 3.75 ± 0.48, n=60) resulted in induction in protrusion formation when compared to controls (Figure 5A-E, 5I-J). Interestingly, in comparison to cells overexpressing HA-dAsap^FL^ alone, a higher number of filopodia were observed in cells coexpressing HA-dAsap^FL^ and Myc-dArf6^FL^ (HA-dAsap^FL^+Myc-dArf6^FL^: # of filopodia, 17.20 ± 0.96, n=123, length in µm: 7.85 ± 0.16, n=148) (Figure 5B, 5F, 5I-J). However, we did not observe any change in filopodia in cells co-expressing dArf6^DN^ and dAsap^FL^ (HA-dAsap^FL^+Myc-dArf6^DN^: # of filopodia, 13.35 ± 1.03, n=117, length in µm: 11.11 ± 0.27, n=286) when compared to cells expressing dAsap^FL^ (HA-dAsap^FL^: # of filopodia, 13.19 ± 0.91, n=117, length in µm: 8.00 ± 0.24, n=143) (Figure 5). In line with the deactivating role of Asap via GTP hydrolysis, the co-expression of dArf6^CA^ in cells overexpressing dAsap^FL^ (HA-dAsap^FL^+Myc-dArf6^CA^: # of filopodia, 6.52 ± 0.60, n=115, length in µm: 8.29 ± 0.41, n=58) led to a reduction in number but not length of filopodia when compared to dAsap^FL^ (HA-dAsap^FL^: # of filopodia, 13.19 ± 0.91, n=117, length in µm: 8.00 ± 0.24, n=143) and dArf6^CA^ (Myc-dArf6^CA^: # of filopodia, 2.65 ± 0.32, n=120, length in µm: 0.42 ± 0.20, n=51) expressed individually (Figure 5B, 5D, 5H-J).

**Figure 5:**
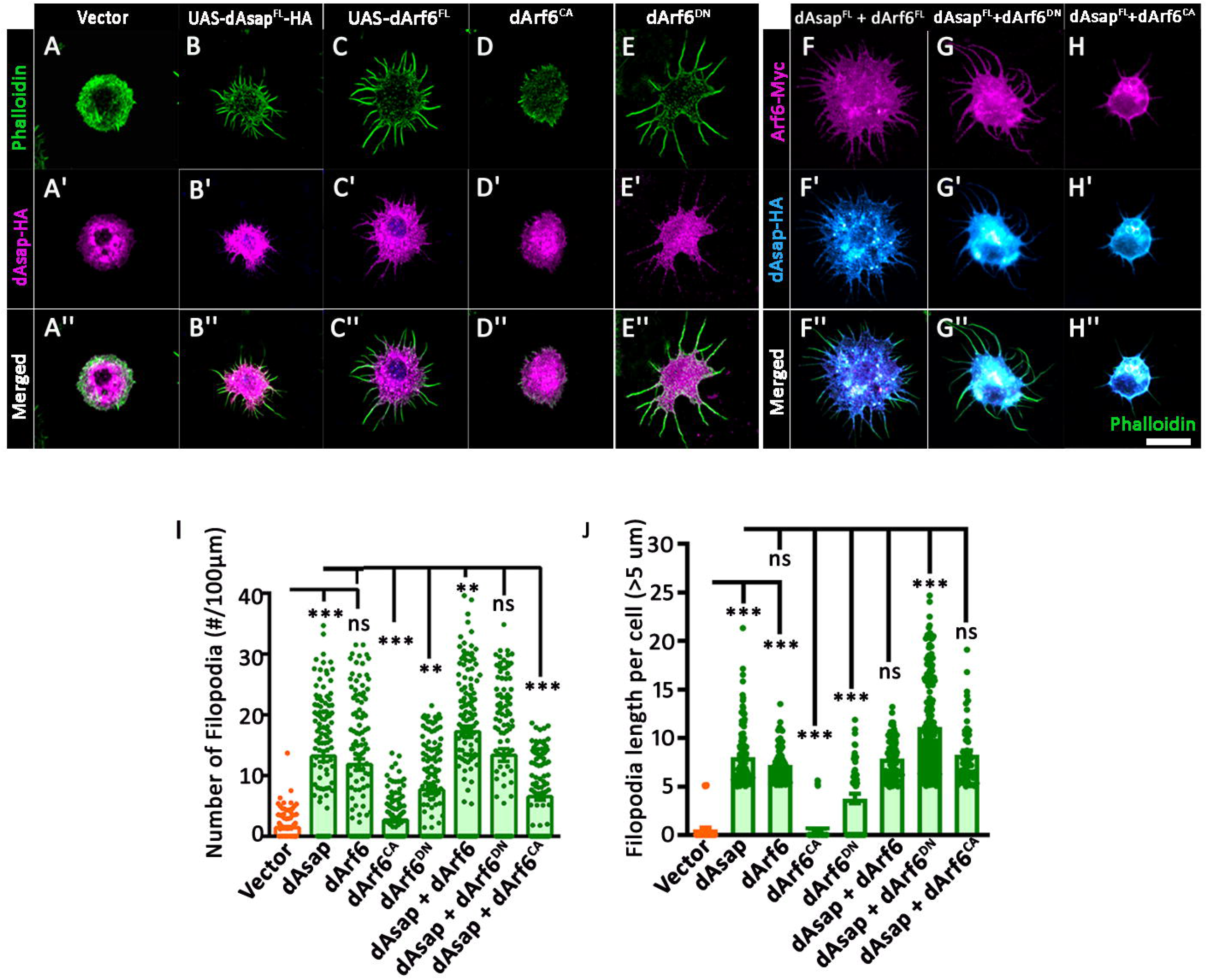
dAsap modulates dArf6 activity to regulate the actin cytoskeleton, forming cellular protrusions in S2R+ cells. **(A, A’’-E, E’’)** Representative images of transfected S2R^+^ cells with empty vector, C-terminal HA tag constructs of dAsap, full length, constitutively active and dominant negative C-terminal Myc tag constructs of dArf6. **(B, B’’, C, C ‘‘)** The overexpression of dAsap or dArf6 resulted in numerous cellular protrusions in S2R^+^ cells. **(D, D’’-E, E’’)** In contrast, overexpression of dArf6^DN^ or dArf6^CA^ resulted in less protrusion formation than dArf6^FL^ in S2R^+^ cells. **(F, F’’- G, G’’)** The co-transfection of HA-tagged dAsap and Myc-dArf6^DN^ or Myc-dAsap^FL^ resulted in more filopodia than dArf6^DN^ or dArf6^FL^ alone. **(H, H’’)** Similarly, the co-expression of HA-tagged dAsap and Myc-dArf6^CA^ resulted in decreased filopodia than dArf6 alone or dAsap alone. S2R^+^ cells were immunostained with phalloidin (green), anti-HA (magenta), and anti-Myc (magenta) antibodies. Scale bar 10 µm. **(I-J)**. The graph quantifies the number and length of filopodia per hundred-micrometer cell circumference. All values here indicate mean ± standard error mean. ***p<0.0001,**p<0.001 and ns, not significant. One-way ANOVA followed by post-hoc Tukey’s test was used for statistical analysis. Each data point or n represents an individual cell used for quantification.

Moreover, our immunostaining results revealed co-localization of dArf6 and dAsap with the actin cytoskeleton, underscoring the potential involvement of dAsap in the regulation of dArf6 activity during actin cytoskeleton reorganization (Figure 5B,5F). These observations indicate that dAsap tightly regulates the active and inactive states of Arf6. Moreover, a balance between these states is necessary for protrusion formations, as supported by the overexpression of dArf6^CA^ in the dAsap background.

## Discussion

Membrane protrusions or lamellipodia formation are essential to cell migration. In this study, we highlight the role of *Drosophila* Arf-GAP, dAsap, in lamellipodia formation. Next, we molecularly characterize the requirement of different dAsap domains for this function. Further, we reinforce the role of actin and its regulatory molecules, Enac and Scar, in lamellipodia formation. Lastly, we demonstrate that dAsap controls the lamellipodia protrusion formation in *Drosophila* S2R+ cells via regulation of dArf6 activity.

Typically, S2R+ cells exhibit a flattened morphology characterized by circumferential lamellipodia containing a cortical band of the actin cytoskeleton (Biyasheva *et al*., 2004; Mallik *et al*., 2023a). Despite lacking directional motility, lamellipodia form near the leading edge of migrating S2R+ cells (Biyasheva et al., 2004). Several proteins, including GTPase activity effectors and proteins containing the BAR domain, have been shown to play a crucial role in membrane protrusion formation (Chen et al., 2020). However, BAR domain-containing proteins, like Cip4, Nwk, IRSp53, and Amphiphysin1, are specifically highlighted in this process (Ahmed *et al*., 2010; Itoh et al., 2005; Peter *et al*., 2004)Ref). This study reveals that dAsap disrupts the dominant morphology of S2R+ cells, forming numerous cellular protrusions. Interestingly, the BAR domain of dAsap is dispensable for this physiological function. These findings contrast the previous studies that demonstrate the role of BAR domains in tubule and long protrusion formation in cells (Itoh *et al*., 2005; Peter *et al*., 2004). To rule out the influence of the Arf-GAP domain over the BAR domain function, we also expressed the BAR domain alone in cells. However, we did not observe any significant change in lamellipodia formation. Here, we conclude that the BAR domain is likely ineffective for membrane tubule or protrusion in *Drosophila* S2R+ cells.

Next, we molecularly characterized the role of individual domains of dAsap in lamellipodia formation by overexpressing different domains in S2R+ cells. Our data suggests that the Arf-GAP domain of dAsap is required for protrusion formation. Surprisingly, the function of the Arf-GAP domain does not depend upon other domains for this process, as the deletion of BAR, Ankyrin, and SH3 domains did not affect the Arf-GAP function in S2R+ cells. Moreover, the Arf-GAP domain construct lacking BAR and SH3 domains was able to hydrolyze Arf1-GTP and Arf6-GTP *in vitro*. These data align with previous biochemical and cell culture studies, which elucidated the involvement of Arf-GAP proteins in diverse cellular functions, including the modulation of the actin cytoskeleton during cell spreading, adhesion, invasion, and migration (Campa and Randazzo, 2008; Ha et al., 2008; Moore et al., 2007). To confirm whether the filopodia induced by the ArfGAP domain of dAsap rely upon actin cytoskeleton structures, we probed S2R+ cells, particularly with SCAR and Ena. While SCAR enhances actin nucleation, Ena regulates the actin filament length and branching (Barzik et al., 2005; Machesky et al., 1999). Our findings indicate that SCAR or Ena initially localized at the cell periphery; however, upon dAsap overexpression, these actin cytoskeleton regulatory proteins redistributed to the filopodial structures at the cell periphery. Previous studies on ASAP1, a mammalian homolog, have implicated its role in focal adhesion assembly, podosome, invadosome formation, and oncogenesis (Furman et al., 2002; Muller *et al*., 2010). Consistent with these findings, we report that dAsap co-localized with actin at the cell periphery and filopodial structures to regulate actin reorganization (Figure 1D-F). Additionally, the re-localization of SCAR and Ena from the cell periphery to the long filopodia upon dAsap overexpression suggests its potential interaction with the actin polymerization machinery in forming these cellular protrusions.

ArfGAP and Arf proteins play crucial roles in the intricate regulation of cellular processes, including filopodia, lamellipodia, or protrusion formation (Dias et al., 2013; Humphreys et al., 2012). The regulatory interplay between ArfGAP and Arf proteins orchestrates the spatiotemporal dynamics of cytoskeletal remodeling and filopodial extensions (Randazzo et al., 2000). The coordinated action of ArfGAP and Arf in regulating filopodia involves intricate signaling cascades, including the modulation of downstream effectors and actin dynamics. Moreover, dAsap, the mammalian homolog ASAP1, has been shown to have GAP activity on Arf proteins that is consequently required to induce membrane ruffles and focal adhesions (Liu et al., 2002; Luo et al., 2007; Randazzo et al., 2000). Owing to these roles of ASAP1, we asked whether the ArfGAP domain of dAsap controls actin-dependent filopodia formation via its GAP activity. Indeed, consistent with *in vitro* investigations on mammalian ASAP1, our data supports that dAsap prefers Arf1 over Arf6 *in vitro* for GAP activity (Kam *et al*., 2000; Luo et al., 2007). Further, our data corroborates with *in vivo* studies in *Drosophila* that highlight the role of dAsap as a GAP for Arf1, maintaining Golgi integrity during cleavage furrow biosynthesis (Rodrigues *et al*., 2016), synaptic morphogenesis (Mallik *et al*., 2023b) and locally inhibiting dArf6 activity during ommatidia patterning in pupal eyes (Johnson *et al*., 2011). Prompted by these results, we investigated whether dAsap relies on dArf1 or dArf6 GTP activity to induce filopodia in S2R+ cells.

Consequently, we demonstrate that the overexpression of dArf6 in S2R+ cells induces the formation of cellular protrusions similar to dAsap overexpression. In contrast, dArf1 overexpression leads to lamellipodia formation in S2R+ cells and actin-polymerization, aligning with prior reports (Humphreys et al., 2012; Rocca et al., 2013). Therefore, we focused on further investigations involving dArf6. Moreover, dAsap, housing an ArfGAP domain, is poised to interact with dArf6, thereby orchestrating the dynamics of the actin cytoskeleton in S2R+ cells. Our investigations reveal the indispensable role of the GAP activity of dAsap in regulating actin polymerization within S2R+ cells, as the absence of the ArfGAP domain reduces protrusions.

Furthermore, co-expression of dAsap and dArf6 demonstrates an augmented actin polymerization, leading to an increased number of filopodia compared to individual expression of dAsap or dArf6. The observed inhibition of protrusion formation in dArf6^CA^ compared to dArf6^FL^ or dArf6^DN^ underscores the crucial requirement for properly recycling Arf6 between its GTP/GDP-bound states for functional efficacy. This notion is supported by previous work indicating that hyperactivation of Arf6 interferes with clathrin-mediated endocytosis, as evidenced by impaired transferrin uptake/internalization upon overexpression of the Arf6GEF, EFA6 (Boulakirba et al., 2014). Likewise, findings in PC12 cells indicate neurite outgrowth inhibition by Arf6^CA^, while Arf6^DN^ or Arf6^FL^ enhances neurite growth, aligning with previous studies (Kobayashi and Fukuda, 2012). Additionally, dArf6^CA^ impedes protrusion formation in dAsap-expressing cells, while dArf6^DN^ shows no significant impact on protrusion formation in S2R+ cells. Thus, the dynamic modulation of dArf6 between its GTP/GDP-bound states emerges as a requisite mechanism for orchestrating the actin cytoskeleton and forming cellular protrusions through dAsap-dependent signaling pathways. While these findings suggest a requirement for both dAsap and dArf6 in actin remodeling, it is essential to note that the intricacies of their functional relevance necessitate further investigation through knockout or knockdown experiments, along with additional cellular assays. These studies will provide a more comprehensive understanding of the roles played by dAsap and dArf6 in actin remodeling processes.

Given that the downstream modulation of actin cytoskeleton dynamics by dAsap and dArf6 may involve various effectors, such as actin-binding proteins and regulators of actin nucleation (Kunda et al., 2003). Furthermore, co-ordinative activity between actin cytoskeleton assembly and Arf6-mediated intracellular trafficking has been implicated in cancer cell migration (Marchesin et al., 2015). Hence, our study on dAsap could shed potential insights into cancer pathology, which needs further investigation.

In conclusion, our study proposes a model in which dAsap overexpression inhibits dArf6 activity, subsequently regulating the actin nucleation pathway for filopodia formation in S2R+ cells (Figure 6). The intricate interplay between dAsap, dArf6, and associated signaling pathways contributes to filopodia formation in S2R+ cells. Further investigations involving the suppression or elimination of endogenous levels of dAsap and dArf6 are warranted to unravel the intricate details of their interaction in regulating protrusion formation in S2R+ cells.

**Figure 6:**
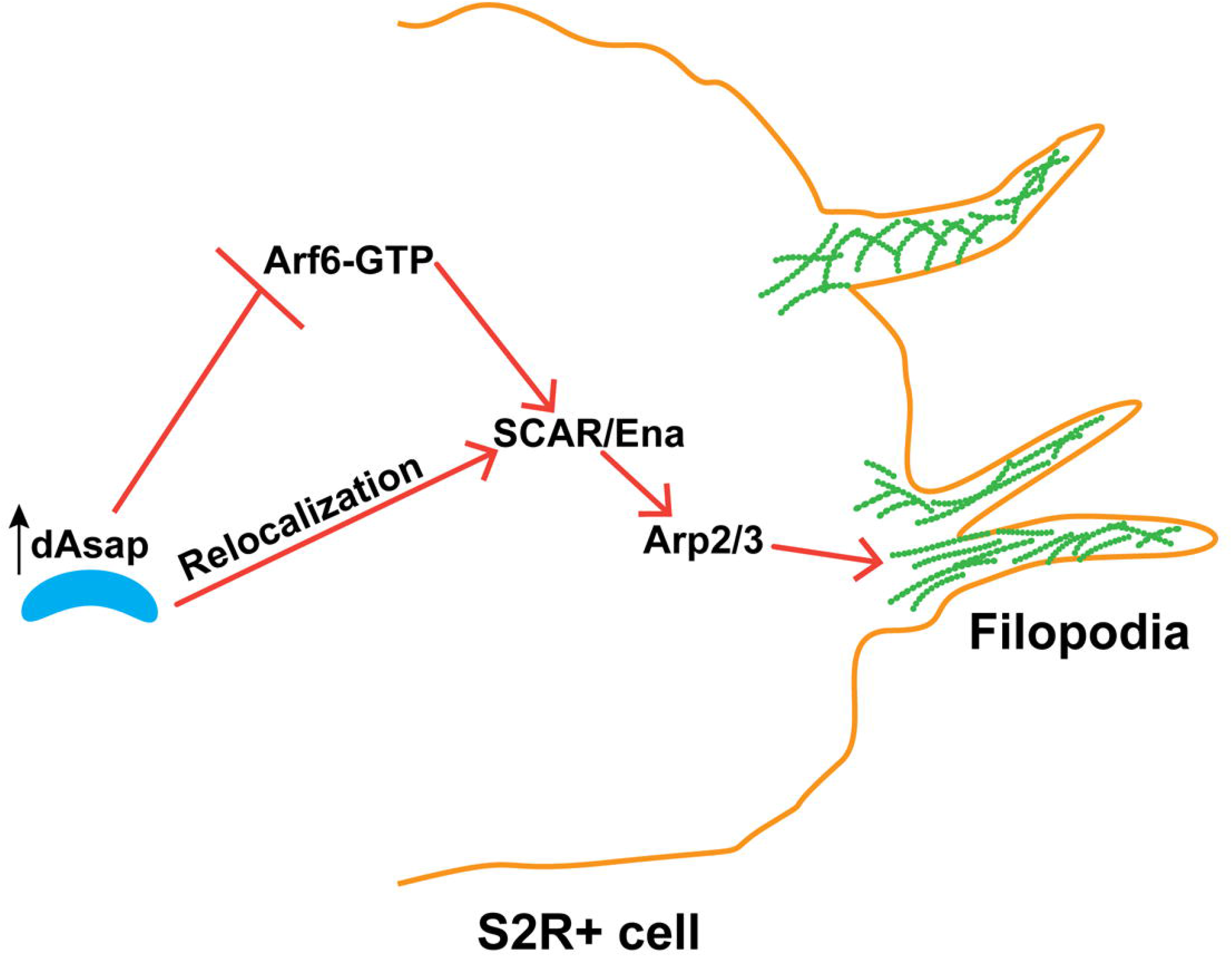
Model depicting filopodia formation through dAsap-mediated Arf6-dependent actin regulatory pathway in S2R+ cells. Based on our findings, we propose this model where upregulation of dAsap inactivates dArf6, further activating the downstream actin polymerization machinery and thus regulating actin nucleation in S2R+ cells. Furthermore, our observations indicate that the over-expression of dAsap induces the re-localization of actin regulatory proteins to the site of filopodia formation. Building upon this concept, we suggest that the regulatory influence of dArf6 induces filopodia formation that depends on GAP domain of dAsap.

## Material and Methods

### *Drosophila* cell culture, transfection, and filopodia quantification

*Drosophila* S2R+ cells were maintained in 1X Schneider’s *Drosophila* media (Thermo Fisher Scientific), supplemented with heat-inactivated 10% FBS, 50 U/ml penicillin, and 50 μg/ml streptomycin. The cell culture was maintained at 25LC (Mallik *et al*., 2023a; Mallik et al., 2017). Transient transfections were performed on approximately 3 × 10^5^ cells using C-terminal HA-tagged constructs of dAsap, including dAsap-pAWH, dAsap^BAR^-pAWH, dAsap^BPZ^-pAWH, dAsap^ΔBAR^-pAWH, dAsap^ΔZ^-pAWH, and dAsap^ΔPZ^-pAWH. Additionally, S2R+ cells were transfected with C-terminal Myc-tagged dArf6 constructs, including dArf6^FL^-pAWM, dArf6^CA^-pAWM, dArf6^DN^-pAWM, dArf1^FL^- pAWM, dArf1^CA^-pAWM and dArf1^DN^-pAWM using the Mirus TransIT transfection agent at a ratio of 2 µl per 1 µg of DNA for single transfection. Peripheral protrusions were quantified by enumerating the number and length of filopodia originating from the cell periphery and normalizing them by the cell perimeter. This calculation provided the number of filopodia per 100 µm and the protrusion length greater than 5 µm for each cell. As described previously, the bar graphs and statistical analysis were made using GraphPad Prism software (Mallik and Frank, 2022; Raut et al., 2017; Sekhar et al., 2019).

### Plasmid constructs

Full-length Asap, Arf1, and Arf6 cDNA were amplified from adult *Drosophila* and cloned into C-terminal HA or Myc-tagged destination vectors pAWH or pAWM, respectively, using the Gateway cloning method. The point mutations were done in dArf6 or dArf1 to form constitutive active (Q67L for Arf6) or (Q71L for Arf1) and dominant negative (T27N for Arf6) or (T31N for Arf1) by using site-directed mutagenesis (SDM kit from NEB) kit, respectively. Next, these point mutants were cloned into C-terminal tagged destination vectors for transient expression in S2R^+^ cells. Similarly, dAsap deletion constructs, dAsap^ΔBAR^, dAsap^ΔPZ^ and dAsap^ΔZ^ were created using SDM and cloned into the *Drosophila* destination vectors. pAWH and pAWM destination vectors were procured from the *Drosophila* Genomic Research Centre (DGRC). The primers for these constructs are included in supplemental table S1.

### Protein expression and purification

Full-length Arf1 and Arf6 were cloned into a pGEX-KG bacterial expression vector for biochemical analysis. The *Escherichia coli* BL21 codon plus cells expressing GST-Arf1 or GST-Arf6 were lysed, affinity purified on glutathione-Sepharose 4B resin column, and digested with Thrombin. The cleaved proteins were eluted and concentrated for further use. The PZA domain of dAsap was cloned in *Pichia pastoris* expression vector pPICZA at KpnI and SacII sites. The His-tagged dAsap^PZA^ protein was expressed and purified following manufacturer guidelines (Invitrogen, USA).

### Malachite green assay

The malachite green substrate is prepared in the ratio of 2:1:1:2 by mixing malachite green (0.08%, w/v in 6N H_2_SO4), ammonium molybdate (5.7% w/v in 6N HCl), polyvinyl alcohol (2.3%, w/v in water with heating) and water (Chang et al., 2008; Rowlands et al., 2004). The assay buffer contains 20 mM HEPES pH 7.5, 150 mM KCl, and 5 mM MgCl_2_. Before starting the reaction, 10 μM of each Arf1 and Arf6 purified protein was exchanged with 1 mM GTP in the presence of 3 mM of DMPC, 5 mM EDTA, and 0.1% sodium cholate in the assay buffer for one hour at room temperature. The reaction followed the increasing concentration of ASAP^PZA^ (P-PH, Z-ArfGAP, A-Ank) protein starting from 0, 0.1, 0.2, 0.5, 1, 2, 5, and 10 μM and incubated at room temperature for 30 minutes. Further, 160 μl of malachite green reagent was added to the 100 μl of Arf1 or Arf6 and ASAP^PZA^ protein mix in a 96-well plate. The reaction was quenched with 20μl of 3.4% sodium citrate after 5 minutes and kept at 37LC for 30 minutes. Next, the absorbance was recorded at 620 nm, and a standard curve was generated using potassium phosphate for every independent experiment.

### Immunofluorescence

For immunofluorescence analysis, S2R+ cells were fixed in 4% formaldehyde for 10 minutes and blocked with 2.5% BSA. Following this, cells were washed with 1X PBS containing 0.1% Triton-X-100 (Mallik et al., 2017). The fixed cells were then incubated overnight with primary antibodies at 4°C, including anti-HA (1:200), anti-Myc (1:200), anti-SCAR (1:100), anti-Ena (5G2, 1:100), and DAPI for nuclear staining. On the following day, after thorough washing, cells were incubated with secondary antibodies for 90 minutes at room temperature. The secondary antibodies used were Alexa Fluor-conjugated Phalloidin 488 (for actin labeling) (1:500), anti-mouse 568, and anti-rabbit 633. For microscopy, S2R+ cells were adhered to Concanavalin A-coated coverslips and imaged using a 63X objective in Apotome and Olympus FV3000. Representative images were captured at a 100X objective with an oil immersion lens in Olympus FV3000.

### Western blot

For western analysis, 1×10^6^ S2R^+^ cells expressing HA-tagged dAsap^FL^ and empty vector were harvested and lysed using lysis buffer (50 mM HEPES [pH 8.0], 100 mM KCl, 2 mM EDTA, 1 mM dithiothreitol [DTT], 1% Nonidet P-40 (NP-40), 2 mM EDTA, 2 mM phenylmethylsulfonyl fluoride and Protease Inhibitor Cocktail) and spun to clear the cell debris. Protein was estimated using the BCA method, and 50µg protein was loaded in each well and separated on 12% SDS gel. Protein was transferred on PVDF membrane (Amersham Hybond) at 4^0^C for 3 hrs, and then it was blocked for 1 hour with 5% skim milk diluted in 1X TBST (Sigma-Aldrich). Overnight incubation was done in primary antibodies SCAR (1:500) and anti-Ena (1:500) at 4L. GAPDH was used as the loading control. The membrane was washed with 1X TBST and again incubated with HRP-conjugated secondary antibodies (1:10000) for 90 minutes. Finally, signals were detected using a Licor bioscience imaging system.

## Acknowledgments

We thank the Developmental Studies Hybridoma Bank at the University of Iowa, USA, for providing the antibodies utilized in this study. Fly stocks were obtained from the Bloomington *Drosophila* Stock Center. We extend our appreciation to the members of the Vimlesh lab for their valuable comments and discussions during this research. We would also like to thank Dr. Sunando Datta, IISER Bhopal and his lab members for useful comments during the preparation of this manuscript. Research fellowships from the Department of Biotechnology (DBT-JRF) partially supported BM and SK, respectively. This phase of the investigation received financial support through grants from the Department of Science and Technology (DBT), India (DBT Project No-BT/PR/26071/GET/119/108/2017), awarded to VK.

## Author contributions

SK, BM and VK designed the research; SK, BM, ZM and AB performed the research; SK, BM, ZM, and VK analyzed the data; BM, SK and VK wrote the paper.

## Declarations of interests

The authors declare no competing financial interests.

## Data availability

The data and reagents used in this study will be shared upon request.

## Notes

### Competing Interest Statement

The authors have declared no competing interest.

