## Supplemental Table 1 for "*Drosophila* Asap regulates cellular protrusions via dArf6-dependent actin regulatory pathway"

**Supplemental information**

| **Sl. No.** | **Name** | **Sequence** | **Remarks** |
| --- | --- | --- | --- |
| 1 | dAsap^FL^ GW FP | CACCATGCCGCCATCCCTGATTGCC | Forward primer for cloning of dAsap^FL^ into the entry vector. |
| 2 | dAsap^FL^ GW RP | ATCAGGCAGCATATGCACGAAACTGA | Reverse primer with stop codon removed for cloning of dAsap^FL^ into the entry vector. |
| 3 | dAsap ^BAR^ GW RP | TACCTTCTGCTTGATCTCATGCAGC | Reverse primer for cloning BAR domain into the entry vector. |
| 4 | dAsap ^BPZ^ GW RP | TCAGTCGCACCGTAAGTCGTTGTCA | Reverse primer for cloning BPZ into the entry vector. |
| 5 | dAsap ^DeltaPZ^ FP | AACGACTTACGGTGCGACTTAGAGC | Forward primer for deleting PZ domain by site-directed mutagenesis. |
| 6 | dAsap ^delta PZA^ RP | GTGGAGGGAGTAACCAGCTCCCCCACT | Reverse primer for deleting PZ domain by using site-directed mutagenesis. |
| 7 | dAsap ^delta BAR^ GW FP | CACCATGGAGAAGCTGCATGAGATCAAGC | For cloning of dAsap fragment with deleted BAR domain region into the entry vector. |
| 8 | dAsap ^delta PZ^ FP | AACGACTTACGGTGCGACTTAGAGC | Forward primer for deleting Z domain by site-directed mutagenesis. |
| 9 | dAsap ^delta ZA^ RP | TTGCAGCTCCACCAGACTTGGACTC | Reverse primer for deleting Z domain by site-directed  mutagenesis. |
| 10 | Ph Kpnl FP | CCGGTACCATGGATTTTGAGCGAGTTGACAAT | Forward primer for cloning of PZA domains into pPICZA. |
| 11 | Ank SacII RP | GGCCGCGGCTCTTAATGGCACATTCAATAAG | Reverse primer for cloning of PZA domains into pPICZA. |
| 12 | dArf1^FL^ BamHI FP | AAGGATCCATGGGAAACGTATTCGCGA | Forward primer for cloning dArf1^FL^ into pGEXKG. |
| 13 | dArf1 HindIII RP | AAAAAGCTTTAGCGATTAGCGTTCTTCAATT | Reverse primer for cloning dArf1^FL^into pGEXKG. |
| 14 | dArf6 BamHI FP | AAGGATCCATGGGAAAGTTACTATCAAAAATTTTC | Forward primer for cloning dArf6^FL^into pGEXKG. |
| 15 | dArf6 HindIII RP | AAAAGCTTTCATAACTTATGGTTCGACGTTAAC | Reverse primer for cloning dArf6^FL^ into pGEXKG. |
| 16 | dArf6 T27N FP | AATACGATTCTGTACAAACTGAAAC | Forward primer for point mutation T27N by site-directed mutagenesis. |
| 17 | dArf6 T27N RP | TTTTCCAGCCGCGTCCAG | Reverse primer for point mutation T27N by site-directed mutagenesis. |
| 18 | dArf6 Q67L FP | CTGGATAAGATTCGACCGCTATGGC | Forward primer for point mutation Q67L by site-directed mutagenesis. |
| 19 | dArf6 Q67L RP | CCCACCGACGTCCCACAC | Reverse primer for point mutation Q67L by site-directed mutagenesis. |
| 20 | dArf6 GW FP | CACCATGGGAAAGTTACTATCAAAAATTTTC | Forward primer for cloning dArf6^FL^ into the entry vector. |
| 21 | dArf6 GW RP | TAACTTATGGTTCGACGTTAACCAAATG | Reverse primer for cloning dArf6^FL^ without stop codon into the entry vector. |
| 22 | dArf1 GW FP | CACCATGGGAAACGTATTCGCGAA | Forward primer for cloning dArf1^FL^ into the entry vector. |
| 23 | dArf1 GW RP | AGCGATTAGCGTTCTTCAATT | Reverse primer for cloning dArf1^FL^ into the entry vector. |
| 24 | dArf1 T31N FP | AACACAATTCTGTACAAACTCAAAT | Forward primer for point mutation T27N by site-directed mutagenesis. |
| 25 | dArf1 T31N RP | TTTACCAGCGGCATCCAA | Reverse primer for point mutation T27N by site-directed mutagenesis. |
| 26 | dArf1 Q71L FP | CTAGACAAAATTCGTCCCCTGTGGA | Forward primer for point mutation T27N by site-directed mutagenesis. |
| 27 | dArf1 Q71L FP | GCCACCCACATCCCACAC | Reverse primer for point mutation T27N by site-directed mutagenesis. |

**Table S1:** The following table describes the list of primers used for cloning the constructs used in this study.
